# Somatic selection of poorly differentiating variant stem cell clones could be a key to human ageing

**DOI:** 10.1101/797530

**Authors:** Walter F. Bodmer, Daniel J. M. Crouch

## Abstract

Any replicating system in which heritable variants with differing replicative potentials can arise is subject to a Darwinian evolutionary process. The continually replicating adult tissue stem cells that control the integrity of many tissues of long-lived, multicellular, complex vertebrate organisms, including humans, constitute such a replicating system. Our suggestion is that somatic selection for mutations (or stable epigenetic changes) that cause an increased rate of adult tissue stem cell proliferation, and their long-term persistence, at the expense of normal differentiation, is a major key to the ageing process. Once an organism has passed the reproductive age, there is no longer any significant counterselection at the organismal level to this inevitable cellular level Darwinian process.

**Author Contributions:** WFB conceived the project. WFB and DJMC wrote the manuscript.

## Introduction

The major features of ageing in complex eukaryotes, notably extensive tissue malfunction as a result of the dysregulation of differentiation, have recently been extensively reviewed (for example, see [1-4]). There is much discussion of the genetic and epigenetic changes that could explain these ageing-associated changes, and the factors, including dietary influences, which could influence the rates at which these changes are produced and how the changes could be ameliorated. Adult tissue stem cells and the extent to which there is somatic selection in their evolution also figure in these discussions [5-7]. There is, however, hardly any mention of the possibility that the degenerative changes in those tissues whose maintenance depends on stem cell tissue turnover could be the result of somatic clonal selection for genetic or epigenetic variants that, to an increasing extent, escape some of the constraints of differentiation and apoptosis and so lead to tissue malfunction.

It is widely accepted that cancer is a somatic evolutionary process in which the steps are genetic, or relatively stable epigenetic, changes, each step successively providing some advantage for the development and outgrowth of the cancer (for example, see [8]). For most cancers, a major step in this evolutionary process is escape from the control over the process by which they differentiate into cells with a programmed limited life span, and the most aggressive cancers are often those that have lost most of their capacity to differentiate. We suggest that cancer can thus be viewed as an extreme form of ageing, with respect to the tissue from which the cancer originates, and that similar violations of differentiation programs by adult stem cells are responsible for many normal ageing phenomena.

Any replicating system in which heritable variants with differing replicative potentials can arise is subject to a Darwinian evolutionary process, and these systems must include the continually replicating adult (i.e., not embryonic, pluripotent) tissue stem cells that control most of the tissue integrity of most relatively long lived, multicellular, complex organisms, including humans. Only the limited number of tissues, such as neurons in the brain, where there is no apparent adult tissue turnover, will not be subject to such a Darwinian selective process. As GC Williams [9] said in his much-quoted, perceptive article on human ageing, “We are not machines that simply wear out, because our parts continually reproduce and refresh themselves throughout our lifetime”. However, as Lopez-Otin et al. [4] said in their comprehensive review of ‘The Hallmarks of Ageing’, “Stem cell exhaustion unfolds as the integrative consequence of multiple types of ageing-associated damages and likely constitutes one of the ultimate culprits of tissue and organismal ageing” (for example, see also [10]). What is missing from this statement is an explanation of *why* all this damage leading to exhaustion should somehow accumulate disproportionately in the stem cells.

We propose that it is Darwinian somatic selection leading to the gradual dysregulation of the stem cell differentiation process and eventually to persistent tissue malfunction that is a major determinant of human ageing and its associated wide range of age-related diseases. This proposal is based on the following assumptions: 1) natural selection at the level of the whole organism effectively stops, or is at least greatly weakened, after reproduction is finished, 2) long term maintenance of most adult vertebrate tissues is undertaken by adult tissue stem cells, but 3) after some period of time, adult tissue stem cells become exhausted [2, 4] and are then replaced by ‘fresh’ stem cells.

Medawar in a key article on ageing, entitled “An unsolved Problem of Biology” [11], said “But cancer and the cardiovascular disease are affections of middle and later life. Most people will already have had their children before the onset of these diseases can influence their candidature for selection. In the post-reproductive period of life, the direct influence of natural selection has been reduced to zero and the principal causes of death to-day lie just beyond its grasp.” This fundamental evolutionary perspective on the problem of ageing has been extended by Kirkwood (see e.g. Kirkwood 2005 [12]) who emphasized that “Somatic maintenance needs only to be good enough to keep the organism in sound physiological condition as long as it has a reasonable chance of survival in the wild.” He thus argued that “The evolutionary explanation of aging rests on the principle that natural selection has sought to optimize the allocation of metabolic resources across core processes like growth, reproduction and maintenance”. This then is the basis for the first assumption underlying our proposal as given above. Given that, now, in most human populations the advances in hygiene, nutrition and medical care have led to an average expectation of life that goes far beyond the usual end of the reproductive period, there is a much longer period of human life when the selective constraints discussed by Kirkwood no longer apply than there is in nearly all other species.

The evolutionary concept for ageing proposed by Medawar, Kirkwood and others, did not specifically take into account the role of adult tissue stem cells in the maintenance of tissues such as the gut and the haemopoietic system. However, Kirkwood and others [13, 14] observed age related changes in the histology of stem cells in the small intestine of the mouse and suggested that apoptosis of damaged cells “may be an important system for long term tissue maintenance”. They suggested that that there was a need to investigate the link between the changes with age at the cellular level and those at the organ and tissue level, which fits in well with our proposed ageing mechanism and its further investigation. Lopez-Otin et al. [4] counted stem cell exhaustion, which they characterized as stem cell decline “due to the integrative consequence of multiple types of damage”, leading “to the decline in the regenerative potential of tissues”, as “one of the most obvious characteristics of aging.” This concept is based on the assumption that adult tissue stem cells, because of their persistent and often high rate of division and production of differentiated cells, eventually accumulate irreparable damage, for example, to their replication and protein quality monitoring machinery and to the maintenance of telomere length. It seems highly improbable that this “exhaustion” only occurs towards the end of life. It is for this reason that there are, as we discuss in more detail later, two levels of adult tissue stem cells. One level is functional in maintaining its relevant tissue and has a relatively high rate of replication while the second has a much slower proliferation rate, being almost quiescent. When a currently fully functional stem cell becomes exhausted, this is recognized by mechanisms such as apoptosis and autophagy or senescence followed by phagocytic removal (see e.g. Munoz-Espin and Serrano [15], Kwon et al. [16], Maiuri et al. [17]). The second level of ‘quiescent’ stem cells, which mostly do not become exhausted because of their much lower level of cellular activity, is poised to replace the fully functional level of stem cells as soon as the need arises, namely when a currently fully functional stem cell reaches exhaustion. This then is the basis for our 2^nd^ and 3^rd^ assumptions, as given earlier, in support of our proposed model.

### Background experimental evidence

It was first shown more than 20 years ago that normal human skin contains large numbers of clones of keratinocytes carrying TP53 mutations [18, 19]. Because these mutations are often due to thymine dimers, it was assumed that they were caused by UV in sunlight on exposed areas of the skin. The fact that the mutations are mostly missense mutations with functional effects strongly implies that the TP53 mutations give the clones a growth advantage, just like that found in many cancers and pre-cancerous growths, including colorectal adenomas. When there is a tumor adjacent to such a clone, it carries a different TP53 mutation, suggesting that in the cancer, the TP53 mutation is not a tumour initiating mutation but provides growth persistence (or immortalization) to the tumour (see e.g., [20]), after other mutations have been selected. Recently, a more extensive study of normal skin biopsies, using DNA sequencing, found evidence for frequent clones carrying a wide range of the sort of driver mutations found in cancers, notably those in NOTCH1, a gene whose function is closely tied to the maintenance of adult stem cells in epithelial tissues [21]. There is similar extensive evidence for somatic clonal selection in hematological cells from older individuals (see e.g. [7, 22-25]).

Weissmann’s group showed that older mice have 5 times as many hematopoietic stem cells (HSCs) in their bone marrow as young or middle aged mice [26], but that these ‘old’ HSCs were much less efficient than ‘young’ HSCs at engrafting the bone marrow of irradiated mice. This clearly suggested somatic selection for clones that had lost much of their differentiating potential and as a result were faster and/or more persistent in their continued growth. More recently, it has become clear that elderly humans without cancers may have only a few dominant HSC clones in their normal bone marrow [27, 28] and that these HSCs often carry mutations in DNMT3A and TET2, genes that are key regulators of DNA methylation, which is the main epigenetic mechanism for gene expression control in differentiated cells. Mutations are also found in genes controlling hematopoiesis and in TP53 [3]. This is clear evidence for somatic selection of dominant clones that have outgrown their contemporaries. A comprehensive analysis of gene expression in highly purified HSCs from young and old mice showed increased promoter methylation associated with differentiation-promoting genes in the latter and decreased methylation with genes associated with HSC maintenance [29]. These results are best explained by somatic selection for methylation changes in the older mice, changes that give rise to HSC clones with higher rates of proliferation at the expense of effective differentiation.

A strikingly similar observation has been made in the fruit fly, *Drosophila melanogaster*, by Biteau et al. [30]. They showed a decline in the ability of intestinal stem cells (ISCs) to differentiate properly in old (40 days) compared with young (3 days) flies, driven by the JNK gene. This decline in differentiating ability is accompanied by an increased rate of ISC proliferation.

An interesting, analogous phenomenon has been described by Andrew Wilkie and his colleagues [31, 32], which they call “selfish selection of spermatogonial mutations”. In normal, especially older, males they find clones of spermatogonial cells, effectively the adult tissue stem cells for sperm production, that carry, for example, mutations in the Kras oncogene that are known to cause severe inherited diseases. These mutations appear to be advantageous for the outgrowth of the spermatogonia that carry them but severely disadvantageous to the individuals who inherit these mutations through fertilisation by mutated sperm. They suggest that this form of selfish selection can explain much of the apparent increase in male germ line mutation rates with age.

### Origins of somatic genetic variation

Mutation in dividing somatic cells is an unavoidable phenomenon. However efficient the DNA repair processes, every cell when it divides will transmit at least a few new mutations to its daughter cells. There is therefore, as might be expected and as we have already discussed, abundant evidence for the existence of somatic genetic variation in a wide variety of human tissues (see Risques and Kennedy [33] for a recent extensive review). Stable DNA methylation changes, which underlie the whole cellular differentiation process, and which can also be the basis for selected gene expression changes in cancers, are an alternative source of genetic variation that is analogous to mutation. Such methylation changes are likely to be very important functional changes at the somatic level as they occur at a higher rate and can directly influence the level of expression of a relevant gene. Recent evidence suggests rates of gene methylation in HSCs increase with age [29].

While extreme differences in the rates of mutation or methylation due to severe germ line mutations in, for example, certain DNA repair genes will be expected to increase the rate of somatic mutations and methylation changes that we suggest underlie the ageing process, the normal background error rates will, we propose, be enough to lead to normal rates of ageing. This is entirely analogous to the fact that normal rates of mutation can lead to tumor development, so that genetic instability is not required for tumour initiation [34, 35].

Mutations can either be neutral, meaning they have no functional effect that influences their survival or rate of proliferation, or be selectively disadvantageous leading to a decreased rate of proliferation or survival, or be selectively advantageous resulting in a higher rate of proliferation and/or survival. Cells with neutral mutations can, just by chance, increase in frequency relative to non-mutant cells. While the probability of this is low for any particular mutation, since there is so much of the DNA that can give rise to mutations, clonal variation can arise from neutral genetic variation, but this variation, by its nature, is generally not likely to lead to any relevant functional change. Variants that are disadvantageous to stem cell proliferation will be selected against and so are not expected to play any role in the ageing process. Even quite small selective disadvantages give rise to exceedingly low probabilities of survival. It is therefore only the variants that favor increased stem cell proliferation that are likely to give rise to functionally affected clones. Even quite small selective advantages give hugely increased chances of survival relative to neutral clones (for the classical population genetics analysis behind these arguments see e.g. Cavalli-Sforza and Bodmer [36, 37]).

### Mitochondrial DNA mutations

There have been many studies on the increase of mitochondrial DNA mutations with age (see e.g. Su et al. [38] for a recent review), with unequivocal evidence of a substantial increase in the number of MtDNA mutations with age in the stem cells of human colonic crypts [39]. Mitochondria have their own DNA (MtDNA), which codes for a limited number of mitochondrial proteins or their sub-units and for some functional RNAs. Each cell usually contains from at least a few hundred to more than a few thousand mitochondria. The mitochondria present in a cell as it divides are presumed to be passed on at random to the two daughter cells, allowing for a fair amount of stochastic variation in the behaviors of new MtDNA mutations. There is no obvious direct relationship between the control of nuclear genomic cell DNA replication and that of the mitochondria. MtDNA mutations in cytochrome-c oxidase leading to its inactivity can be readily stained for but only if a reasonably high proportion of the mitochondria in a cell carry the mutation. This has been used for clonal analysis of both large and small intestinal crypts and there is no clear evidence that lack of the enzyme activity in all the mitochondria of a cell affects its replication potential. There is clear evidence that the mutation rate of the MtDNA is substantially higher than that of the nuclear genomic DNA. The overall features of the MtDNA turnover and the arguments over the effects of MtDNA mutations on a cell’s function even when a high proportion of the mitochondria in a cell carry a functionally effective mutation make it, in our view, very hard to interpret the functional significance of the increase in MtDNA mutations in stem cells with age. It is even possible that mutations that are apparently deleterious to the cell may sometimes provide a ‘selfish’ advantage to the independently dividing mitochondria. Perhaps even this possibility could increase with age beyond the reproductive period, so long as the effect on the cell was not too deleterious.

### Mechanisms of somatic selection in adult tissue stem cells

The key to our model for ageing is not just to suggest an effect of selection on the ageing adult tissue stem cells (see, e.g., Beerman et al. [7]), but to propose the underlying basis for the mechanism of action of the selection and the reason for its detrimental effect on adult tissue functions

Thus, we suggest that the common feature in all the examples of age-related functional changes in adult tissue stem cells that we have discussed is somatic selection for mutations (or stable epigenetic changes) that give rise to an increased rate of proliferation and persistence of adult tissue stem cells at the expense of normal functional differentiation and, eventually, apoptosis or senescence. It is this process that leads to long term tissue malfunction. Once an organism has passed the reproductive age, by which is meant the age range during which they have offspring, there is little, if any, counter selection at the organismal level to this inevitable selfish Darwinian cellular level process that leads to the gradual erosion of the differentiating capacity of adult tissue stem cells through selection for increased stem cell proliferation [12]. While there may be some such phenomena before and during the end of the reproductive age they are clearly most likely to be of much lesser effect. Weaker selective effects at the organismal level, mediated through the benefits that individuals are able to bestow on their younger relatives by helping to care for them [40, 41], may help to explain the relative rarity of tumorigenesis as compared to the normal ageing process during post-reproductive life.

The cancer model of Tomlinson and Bodmer [42], on which we base our ageing model, uses the known cellular composition of the colon as its biological framework. This is based on epithelial stem cells at the base of the colonic crypts that line the surface of the colon. These stem cells control a crypt’s complete turnover about every 4-6 days and give rise to transit amplifying cells, which differentiate into terminally differentiated cells that die by apoptosis and are removed into the lumen of the intestine. The mathematical model assumes that the overall proliferation rate of stem cells can be represented simply by *α* = *α*_3_ − *α*_2_ − *α*_1_, where *α*_3_ is the rate of cell division, *α*_2_ is the rate at which daughter cells become differentiated and *α*_1_ the rate of apoptosis. In a normal steady state *α* = 0, and so, following this model, if non-mutant stem cells have a negligible apoptosis rate *α*_1_ = 0, then *α*_3_ = *α*_2_. This can either be achieved by asymmetric cell division, where when a stem cell divides, one daughter always remains a stem cell while the other always becomes a transit amplifying cell destined to differentiate, or if, as now seems more likely, stem cells on average produce one transit amplifying cell and one stem cell but any given division may produce two daughter stem cells or two transit amplifying daughter cells. In that case the rate at which stem cells produce two daughter stem cells must equal the rate at which they produce two transit amplifying cells. The steady state can then be perturbed by a mutation that decreases the rate at which two transit amplifying cells are produced, so that more daughter stem cells are produced on average than daughter transit amplifying cells. In that case *α*_3_ > *α*_2_ and so *α* > 0 and the stem cells have an increased proliferation rate relative to the steady state maintained by non-mutant stem cells. That is the basis for assuming that mutations which decrease the rate at which a stem cell produces daughter transit amplifying, and then differentiated, cells may be the basis for selection for increased rates of proliferation of the stem cells, at the expense of normal rates of differentiation. This then disturbs the normal steady state ratio between stem and differentiated cells and so is a basis for tissue malfunction.

Why should these effects be concentrated in adult stem cells, as opposed to transit amplifying or differentiated cells? Transit amplifying cells complete a limited number of divisions before differentiating and then undergoing apoptosis and they will therefore require a major restructuring of their cellular growth pathways to reactivate a proliferative phenotype. Although this phenotype would confer a selective advantage at the cellular level, it would almost surely not be reached without several mutations. In contrast, in the adult stem cell, one mutation could be sufficient to alter the rate of proliferation versus differentiation.

The phenomenon of senescence was first described in human *in vitro* cultured fibroblasts but then shown to occur in other cell types as well and, in particular, in pre-cancerous states. Although at one time thought to be a key to understanding the ageing process at the cellular level, it is now realized to be analogous to apoptosis in being a mechanism for recognition of intra-cellular damage (see e.g. Munoz-Espin and Serrano [15]). In most cases, cellular senescence induces a secretory pathway that leads to the phagocytosis of the senescent cell and so, for some tissue stem cells, such as probably for mesenchymal fibroblastic cells, it is an alternative to apoptosis. Thus, for the purpose of our model, senescence can simply be considered an alternative to apoptosis.

Transit amplifying or differentiated cell populations could, in principle, gain a selective advantage through the acquisition of mutations causing delayed apoptosis, but these mutations could only be passed on to a limited number of daughter cells (or none, in the case of the non-dividing differentiated cells), and would thus have only short-term effects.

Mutations occurring in adult stem cells, *when expressing reduced apoptosis only in their descendant differentiated cells*, would produce a continuing stream of differentiated progeny with longer than normal life spans. These cells would possess an advantage over those undergoing normally programmed cell death, and Tomlinson and Bodmer [42] showed that the longer life spans are not consistent with exponential (i.e. malignant cancerous) growth. Thus, when the apoptosis rate in differentiated cells is small relative to the rate of differentiation, the expected result is a relatively large number of extraneous, differentiated cells, effectively an increasingly large benign tumour. In theory then, this process could be a mechanism for ageing; however, it is unlikely to contribute significantly, as the underlying mutation is not expected to spread beyond a single adult stem cell or provide any direct selective advantage to the stem cell that carries it. Stem cells will themselves eventually become ‘exhausted’ and apoptose, as already discussed, and so any descendant population of differentiated cells is likely to be entirely replaced by one undergoing normal programmed cell death. In any case, it seems unlikely that there are common mutations with effects on apoptosis that are large enough to produce sizeable, persistent benign growths emanating from individual stem cell clones.

These arguments against a role for mutations that affect only the function of transit amplifying or differentiated cells lead naturally to the suggestion that the predominant cause of ageing lies in the adult stem cells *themselves* evolving a resistance to apoptosis or, more likely initially, to differentiation. This resistance to differentiation, leading to increased proliferation, will result in greater numbers of mutant stem cells, as explained above, and will confer a selective advantage on them. Johnston et al. [43, 44], developing the model of Tomlinson and Bodmer [42] by incorporating homeostatic feedback, showed that successive genetic or stable epigenetically based changes that either decrease apoptosis or decrease differentiation rates in stem cells lead to a series of increasingly large, stable benign tumours, with increasingly large numbers of stem cells and differentiated cells, the latter still having their life spans limited by apoptosis. The incorporation of feedback enables homeostatic regulation of the stem cells and buffering against stochastic physiological variation which is not possible under the original Tomlinson and Bodmer model. This enables the stem cell number to vary around a fixed mean over time, leading to corresponding minor variation in the balance between stem cell division and differentiation. Cancers, resulting from exponential, continuous, growth only arise after the cumulative effect of a number of such steps exceeds the limitations of benign growth. In this sense, the ageing adult tissue stem cells give rise to the analogue of an adenoma rather than a carcinoma. Decreases in the rates of differentiation, and concomitant increases in the rates of stem cell proliferation, that either preserve or do not greatly change the total numbers of cells present in a region of tissue and so maintain basic functionality, will confer smaller selective disadvantages to the organism as a whole than those which cause an overall large growth in numbers and greater disruption of tissue differentiation, as happens in cancers.

As apoptosis occurs infrequently in normally functioning stem cells and is presumed to occur mainly in response to signs of stem cell exhaustion, failure to differentiate properly is very likely to be the candidate for the selective effect of initial mutations. Resistance to apoptosis will confer no significant selective advantage to cells that are not yet displaying signs of exhaustion. For the *eventual persistence* of the resulting abnormally functioning tissue, there must, however, be further selection specifically against apoptosis of the stem cells carrying the mutations that lead to defective differentiation. This further selection will ensure against full replacement of the overgrown population of abnormally differentiated mutant or epigenetically altered cells by fully functional, normally differentiated tissue derived from a fresh stem cell, originating from an underlying pool of potential fully functional stem cells, as discussed earlier. Such replacement will occur in response to signs of exhaustion, including a failure to differentiate at the normal healthy rate. This can explain the presence of, for example, TP53 mutations promoting immortality in ageing adult stem cells, analogous to the presence of TP53 mutations in late larger colorectal adenomas [45]. Failure of stem cell differentiation, associated with persistent increased proliferation, thus acts as a strongly positively selected phenotype at the level of the stem cell, and a weakly negatively selected phenotype at the level of the organism. It is just such changes that we propose are the candidates for many of the causes of normal ageing.

In contrast to the ageing situation, during life up to the end of the reproductive period there is strong selection at the higher level of the individual whole organism, for the integrity of adult tissues over many cycles of stem cell proliferation. As a result, there are mechanisms that differentially protect the adult tissue stem cell from accumulating various forms of damage, including DNA damage, and that ensure the replacement of a stem cell once it has exceeded a certain level of ‘exhaustion’ (for example, see [10]). One feature of this protective process is, as already mentioned, the existence of two levels of adult tissue stem cells. One level is functional at any given time and is responsible for the clonality of individual colorectal crypts [46], while the second has a slower proliferation rate (some say that cells at this level are quiescent, [47]). The second level ‘quiescent’ stem cells are poised to replace the first level stem cells as soon as the need arises, namely when a currently fully functional stem cell reaches exhaustion. There is direct evidence for these two levels of stem cells in the human gut [48-51]. It is the counteraction of the exhaustion of individual stem cells, presumably by countering apoptosis, that gives rise to the persistence of dysregulated differentiated tissue, as discussed above. With this background, we will now discuss in more detail how the model of Johnston et al. [43], with some modification can, we suggest, explain how selection works on adult tissue stem cells. The aim of such a model is not so much to provide a basis for estimating relevant parameters, such as division and differentiation rates, but to provide a quantitatively and biologically plausible mathematical description of how such selection could work in a way that is consistent with the biological observations we have reviewed.

### Quantitative model

An extension to the model of Johnston et al. [43], which can explain how long term persistence of individual cells can occur and why this is only likely to happen to cells that have already acquired a failure to differentiate at the full healthy rate, is shown schematically in Figure 1. There are two key additional assumptions. The first is that the homeostatic feedback on the number of cells is non-linear and can thus be overwhelmed when the number of cells passes a certain point. The second is the addition of the near-quiescent level of stem cells that can replace exhausted functionally active stem cells. The figure shows a graphical depiction of the differentiation process and its relationship to the various parameters. We retain the same general notation as in the original Johnston et al. [43] model, which relates specifically to cell types within the colonic crypt, as already discussed. We represent the number of quiescent or ‘second layer’ stem cells as *N*_0_, the number of functional stem cells as *N*_1_, the number of semi-differentiated still dividing cells as *N*_2_ and the number of fully differentiated cells as *N*_3_. For each stem cell type we use the parameters *α*_1_, *α*_2_ and *α*_3_ (for quiescent stem cells) and *β*_1_, *β*_2_ and *β*_3_ (for functional stem cells) to represent the apoptosis, differentiation and renewal rates respectively. For transit amplifying or semi-differentiated cells we use the equivalent parameters *δ*_1_, *δ*_2_ and *δ*_3_, and for fully differentiated cells we define a single parameter *γ*, representing their apoptosis rate.

**Figure 1.**
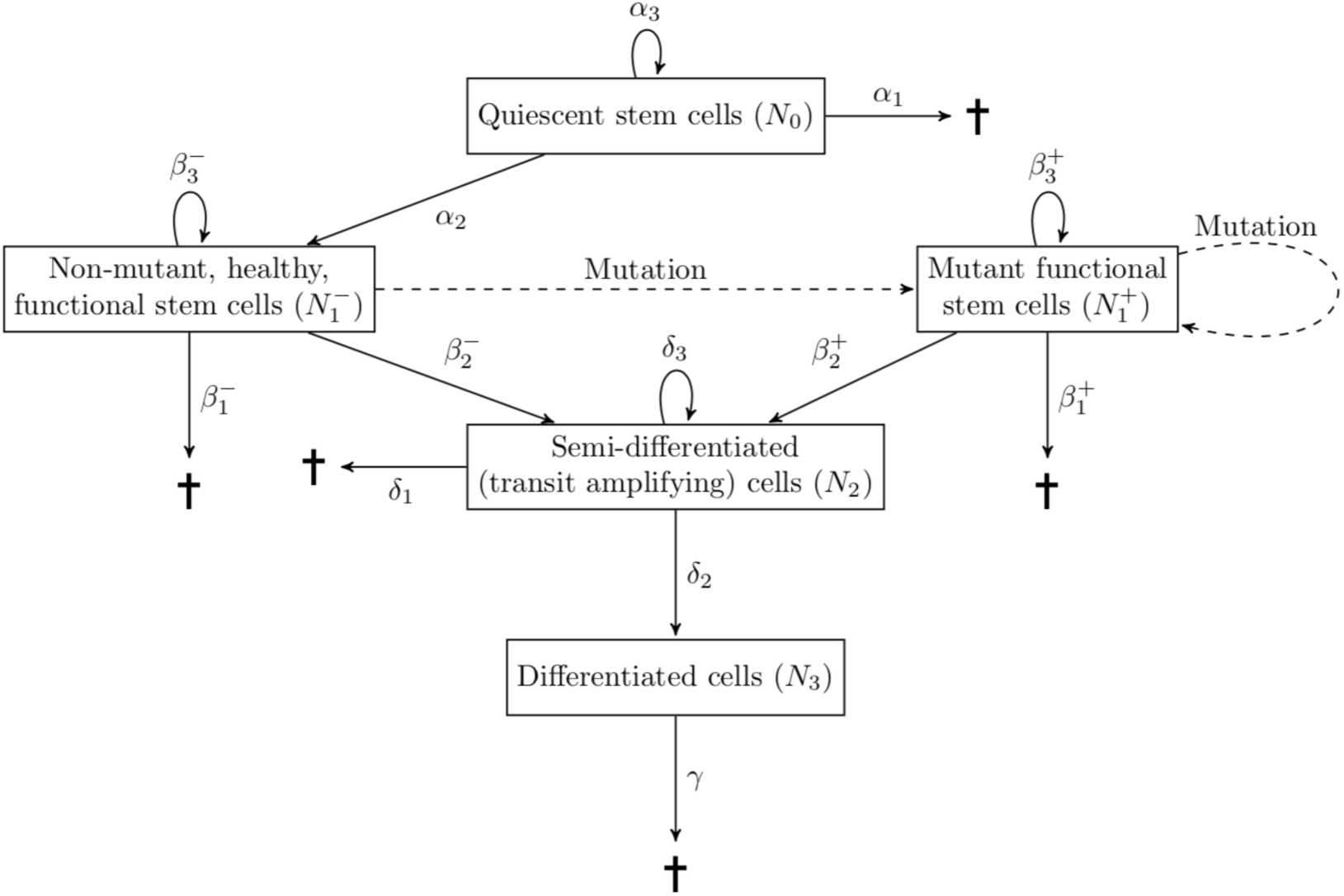
Schematic representation of a model for stem cell proliferation and differentiation. Crucifix signs indicate cell death.

Thus, as depicted in Figure 1, the individual cells of the quiescent stem cell population can, when they divide, die (*α*_1_), become a healthy (non-mutant) functional stem cell (*α*_2_) or self-renew (*α*_3_). Similarly, the healthy functional stem cell can die (*β*_1_), become a transit amplifying semi-differentiated cell, which can self-renew and then become fully differentiated (*β*_2_), or renew (*β*_3_). The same applies to the semi-differentiated cells (*δ*_1_ for dying, *δ*_2_ for differentiating and *δ*_3_ for self-renewal), while differentiated cells can only die by apoptosis or, as discussed above for some tissues, after entering cellular senescence (*γ*).

We assume that when the population of stem or semi-differentiated cells increases, the rate at which new cells are produced also increases, but instead of assuming a linear dependence of per-capita rate on population size we assume that there is a maximum per-capita rate represented by feedback terms in the following equations. This feedback is an essential feature of our model, as it is required to obtain stable states of the relative proportions of the various cell types, where the change in these proportions depends on the magnitude of the effects of mutations on the properties of the functional stem cells. A simple model with no feedback control necessarily results in exponential growth of adult tissue stem cells and so cannot explain what happens with ageing other than giving rise to a cancer. The rates of change of the various cell type counts are as specified by Johnston et al. [43] using the following differential equations:

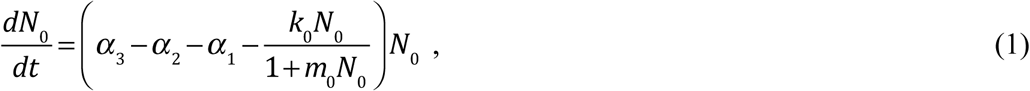

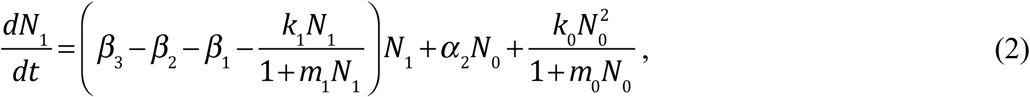

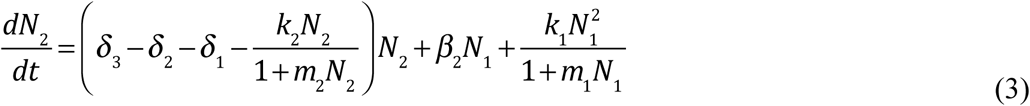

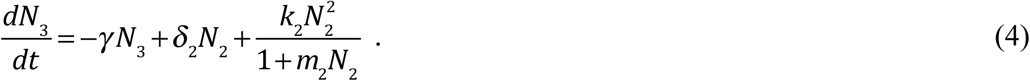

In these equations, the non-linear homeostatic feedback to control cell numbers is captured by the terms 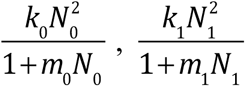, and 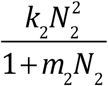. These effectively impose carrying capacities on the three types of cell, which can be overwhelmed when the cells replicate too rapidly. If *m*_0_ = *m*_1_ = *m*_2_ = 0 feedback is linear. Cell numbers beyond the limit imposed by feedback are forced to differentiate into the downstream cell type.

Non-linearity is essential to capture the ability of cells to overwhelm homeostatic control, as this is necessary to produce exponential cancerous growths, as shown by Johnston et al. [43].

The model can be expanded to allow for continual replacement of mutant (+) dysregulated stem cells in the functional layer from a supply of ‘fresh’ non-mutant (-) stem cells produced by the quiescent level of stem cells. We can then show (see Appendix A) that the equilibrium for the total non-quiescent stem cell population size, when the mutant is able to spread, is

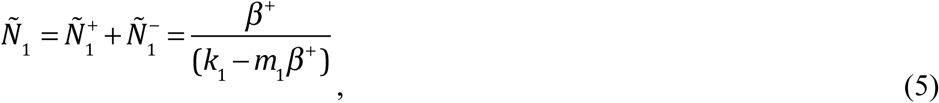

where 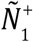 is the mutant stem cell population size, 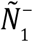 is the non-mutant stem cell population size and *β*^+^ the net proliferation rate of the mutant stem cell population (see Appendix C for the equilibrium value when the mutant does not spread due to insufficient proliferation). This result, surprisingly, does not depend on any properties specific to the non-mutant functional stem cells.

However, if the mutant survives, this does not cause the elimination of the non-mutant type, as it can also be shown (Appendix A) that when growth is not exponential (namely not cancerous), the healthy non-mutant stem cell type is maintained through continual replenishment from the quiescent stem cells. The numbers of non-mutant and mutant cells at equilibrium (Appendix A) are:

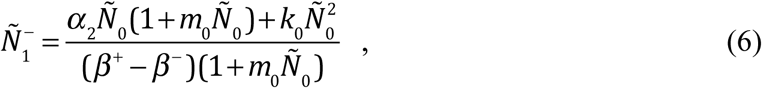

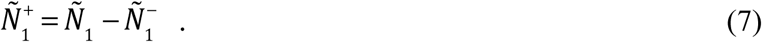

When a new mutant is introduced into a population consisting of an existing mutant type and a healthy non-mutant type, and if it can increase in frequency, the new mutant will entirely replace the original mutant (Appendix A). The new mutation could either be a different mutation in the non-mutant stem cell population 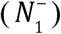 or an additional mutation in the already mutated stem cell population 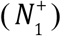.

The effects of mutations on tissue function will be intrinsic to the mechanisms by which mutations affect differentiation and will also depend on the increased number of resulting differentiated and semi-differentiated cells, *N*_3_ and *N*_2_, which themselves depend on 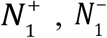 and *N*_0_, the numbers of mutant and non-mutant functional stem cells and quiescent stem cells respectively. A numerical example of an outcome of the model based on the above equations is shown in Figure 2a. This shows equilibrium values for 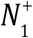 and 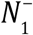 after a series of mutations changing the differentiation 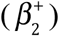 and apoptosis 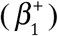 rate parameters in the functional stem cell population. Consistent with our proposed model, the first three mutations confer a reduced differentiation phenotype, while the fourth and final mutation confers almost complete resistance to apoptosis, in effect immortalisation. The numbers of semi-differentiated and differentiated cells descending from mutant and non-mutant stem cells are also shown. Equilibrium states are calculated from modified equations based on Johnston et al. [44] and are given in Appendix C. The final equilibrium state shows that, now, most of the cells are mutant, with a large number of presumably poorly functioning differentiated cells.

**Figure 2.**
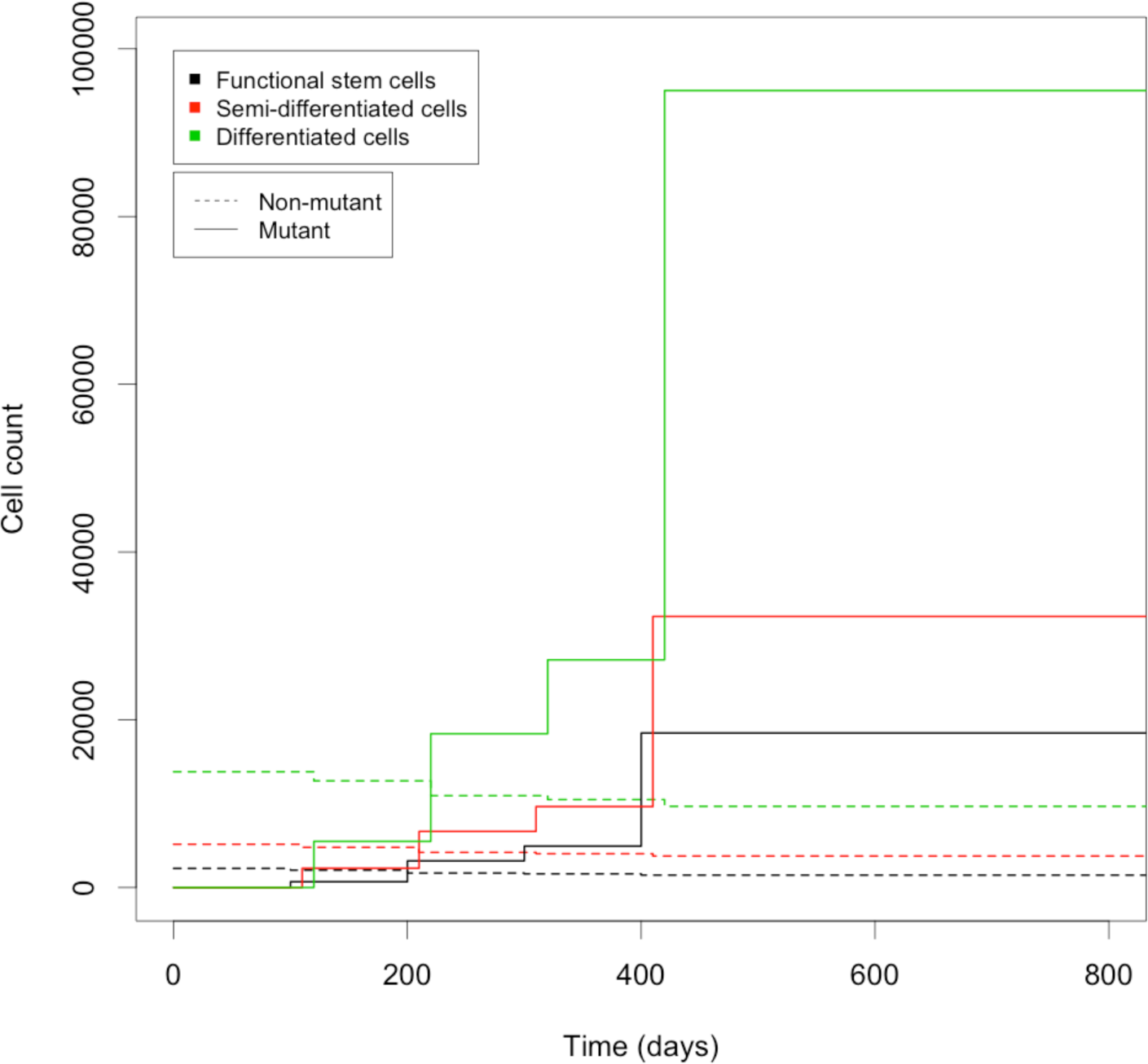
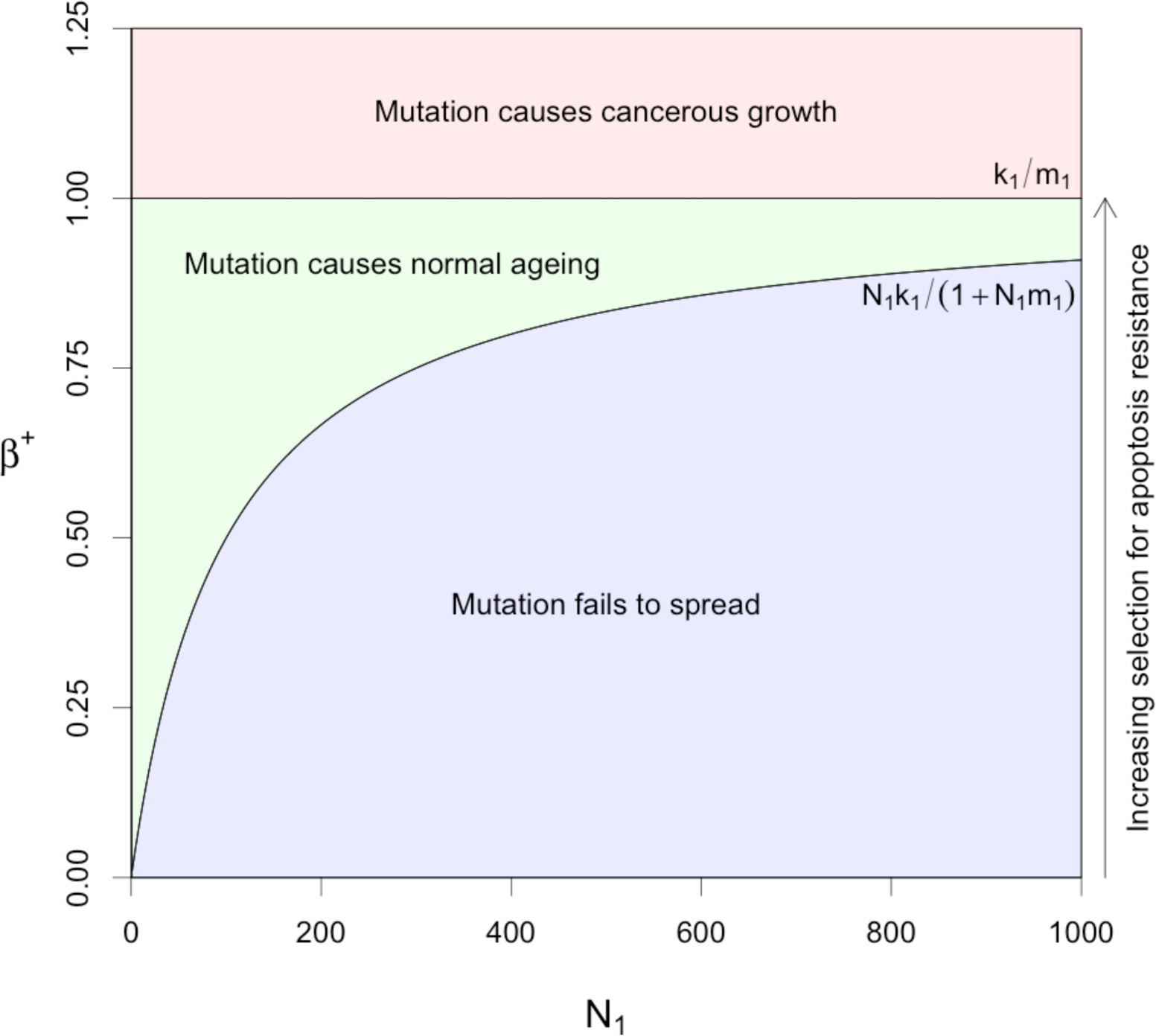
a) An illustrative example of the outcome of the model based on equations 1-4 for a sequence of functional stem cell mutations (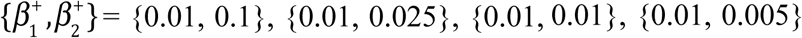, and {0, 0.005}) occurring successively every 100 days. The initial values are those for the non-mutant rates, 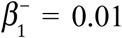 and 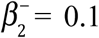, which do not vary. Day zero is a point after the end of the organism’s reproductive lifespan, when selection for healthy tissue functioning has been greatly diminished. Each vertical line marks a new mutation. Its effect on the numbers of each cell type is given by the horizontal lines, which show stable equilibrium cell counts for each new mutation (See Appendix C). Remaining parameters were set to *k*_0_ = 0.1, *m*_0_ = 0.1, *α*_1_ = 0.01, *α*_2_ = 0.05, *α*_3_ = 1, *k*_1_ = 0.01, *m*_1_ = 0.01, 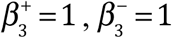, *k*_2_ = 0.001, *m*_2_ = 0.001, *δ*_1_ = 0.1, *δ*_2_ = 0.5, *δ*_3_ = 1 and *γ* = 0.5 and did not vary, on the assumption that the mutations only affected the parameters of the functional stem cell population, *N*_1_. b) Parameter space leading to normal ageing effects (stable growth) and cancerous effects (exponential growth) versus functional stem cell population size. Parameters *k*_1_ and *m*_1_ were both set to 0.01.

Regions of stability in the *β* parameter space for the system with saturating feedback in the functional stem cell population are shown in Figure 2b, plotted against the functional stem cell population size, *N*_1_. If *β*^+^ < *N*_1_*k*_1_ /(1+ *N*_1_*m*_1_), then the mutant stem cells cannot sustain their number and the mutant contribution to the tissue region becomes extinct (see Appendix A), resulting in a non-mutant healthy stable tissue region (blue region). If *β*^+^ > *k*_1_ /*m*_1_ the growth saturation limit is exceeded, and the cell populations grow with no bound as in a cancer [43, 44] (red region). The green region corresponds to normal ageing, where mutant functional stem cells can sustain their growth but do not overwhelm homeostatic non-linear feedback so as to grow exponentially and turn into a cancer. This prediction that the normal aging space shrinks as the cell population expands suggests that the normal aging process should actually decay within a single tissue region, after an initial rapid phase, once post-reproductive life ends, rather than accelerate. Once overgrowth of poorly differentiating cells has occurred, scope for further growth becomes more and more limited without it becoming cancerous, which would be selected against more strongly at the level of the organism.

## Discussion

We have shown how adult tissue stem cell growth based on selection, after the end of the reproductive period, for increased stem cell proliferation at the cost of reduced differentiation and eventually elimination of apoptosis, can lead to immortalisation of aberrantly differentiating adult tissue stem cells and stable malfunctioning mature tissue but not cancer. We suggest that this can explain the ageing process, particularly in humans, because the current expectation of human life is so far extended beyond the end of the reproductive period. The model predicts reduced efficacy of the adult tissue stem cells, for those tissues that are continually turning over. This is compatible with observations of increased numbers of somatic clones with mutations that are often found in cancers, as a function of increasing age. In the particular case of HSCs this also fits with the evidence of changes in their patterns of differentiation associated with increasing age, and also associated with oligoclonality of the HSCs.

Our quantitative model is a key part of our case for the role of somatic selection against differentiation and apoptosis in adult tissue stems cells as a major feature of the ageing process. Essential features of the model are that it includes the minimal number of plausible cell compartments that is known to model adult tissues, using crypts in the normal colon as our biological example, with the minimum necessary number of associated parameters, and that stable states cannot be established without some form of feedback. The biological plausibility of the model, in relation to the observations we have reviewed, is most directly demonstrated by Figure 2b, which shows the limited range of values of *β*^+^, the net functional stem cell reproduction rate, in relation to *N*_1_, the number of functional stem cells, that can lead to functional differentiated tissue without giving rise to a cancer.

The quantitative implementation of the model confirms that failure of functional adult stems cells to differentiate at the normal rate increases their rate of renewal, and so leads generally to stable overgrowths of cells. The model, however, only leads to a cancer when differentiation rates reduce to below a certain threshold. Within growths associated with mutations, or stable epigenetic expression changes, that increase the net functional stem cell rate of proliferation, *β*^+^, the proportions of the various cell types are different to those observed within healthy tissue, exhibiting an excess of semi-differentiated and differentiated cells. These predictions are entirely consistent with observations of aged tissue.

Clearly for those tissues where there is no apparent turnover connected with tissue stem cells, for example neurons in the brain, ageing may be mostly due to wear and tear with age connected with constant use, just like the parts of a car. However even such tissues are likely to be materially affected by declining function of the stem cells for supporting tissues and cell types, for example, the glial cells in the brain. As most tissues are subject to significant cellular turnover, we conclude that our model is likely to explain many of the features of human ageing. Although such tissues are also likely to experience wear and tear like any other, stem cell turnover ameliorates these effects via the replacement of damaged cells, and this is almost certainly one of its explicit functions. Therefore, in tissues and cell types where there is an adult stem cell turnover system, we predict that Darwinian natural selection of mutant stem cells will be responsible for the majority of age-associated damage.

Spatial expansion of mutant clones at the expensive of other cells has been shown to occur in various tissues [21, 52-54]. Although growths of poorly differentiating stem cells descending from a single mutant clone are unlikely to account for the aging of an entire tissue, the rate of successful selective increase of such mutations becomes much higher after the end of the reproductive life of the organism. Under such circumstances, the relevant mutations will occur and spread multiple times in different tissue regions, thereby accounting for significant damage. This would, for example, explain how liver spots are an indicator of aging skin, but occur in relatively high numbers, with each only moderate in size compared to the entire surface of the skin. A mutation causing a growth which took over a very large part of a tissue would most likely be one overwhelming any kind of homeostatic feedback and therefore, in effect, be cancerous, as our model shows (Figure 2b).

We have emphasized the end of reproductive life as a stage when selection is greatly relaxed at the level of the organism, following Medawar and Kirkwood. However, it is important to point out that the reproductive value of an organism, defined by R.A. Fisher as the extent to which “persons of this age, on the average, contribute to the ancestry of future generations” [55], usually falls gradually after the start of the reproductive period in adolescence, implying that selective pressure for an organism’s integrity will relax slowly over its entire adult life. This is largely because the chance of death due to unpredictable events, such as infectious disease, starvation or attacks by predators, increases with time. However, after reproductive life ends, an individual’s reproductive value consists solely of the aid it can provide to its relatives [40, 41], and so selection will be especially diminished after this point. Whilst human males have no strict end to their reproductive lifespan, reproductive behaviour on average tales off greatly by late middle age to the point that it can effectively considered to be over.

On the assumption that, as we have argued, the somatic selection of variant clones with dysregulated differentiation is inevitable and may be a key to human ageing, the basic scientific problem is the lack of understanding of the normal processes of tissue homeostasis before the onset of organismal ageing, and of the changes in these processes with organismal ageing. A systematic study of stem cells and their immediate progeny in a variety of tissues and at different organismal ages, analogous to the study of HSCs by Xie et al. [28], should help to reveal the changes that are associated with ageing. Similar comparisons could also be made between healthy individuals and those with premature ageing, such as Werner Syndrome patients. Analyses of mRNA in single cells should reveal the balance of different cell types at different stages of differentiation, while direct DNA and methylation sequencing should reveal the genes in which there are somatic mutations or stable methylation changes in expression in older organisms compared to younger organisms. It will then be interesting to see to what extent the genes in which there are significant changes correspond to the genes in which there are driver mutations or stable methylation changes in cancers from the relevant tissue, as is the case for the somatic mutations found in HSCs from normal healthy elderly individuals [28]. In agreement with this possibility, Slack et al. [56] have suggested that drugs that attack some of the cancer promoting pathways, for example, those associated with mutations in Kras and PTEN, might counter ageing effects. This idea is consistent with the notion that abnormally functioning adult stem cell clones may accumulate genetic or epigenetic changes by a process of somatic selection for driver mutations or stable methylation changes that is similar to that found in cancers. Our model would also predict that, in addition to possessing relevant epigenetic changes and DNA mutations in adult tissue stem cells, aged individuals would lack fully functional homeostatic control of stem cell populations, although experimental investigation of this is likely to be challenging.

Any general ‘anti-ageing’ treatment must counter successfully all, or at least many, of the Darwinian selection processes acting against normal homeostasis of all the different vital tissues of the ageing human body that are maintained by adult tissue turnover. That is a considerable challenge that may never be fully met. In the meantime, the obvious priority is to continue to try and ameliorate the diseases of old age, including cancer, rheumatoid and osteoarthritis, autoimmune disease and immune function decline, Parkinson’s disease, dementias, strokes, heart disease and type 2 diabetes. In this way, we can at least hope to continue to improve the quality of life of the elderly within our existing maximum life span of approximately 100 years.

## Acknowledgements

This work was supported by JDRF Grant 5-SRA-2015-130-A-N and Wellcome Trust Grants 107212/Z/15/Z and 203131/Z/16/Z.

## Appendix

### A) Equilibrium cell count and mutant frequency for functional stem cells

The equations for the increase of mutant 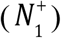 and non-mutant 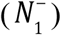 functional stem cell counts are:

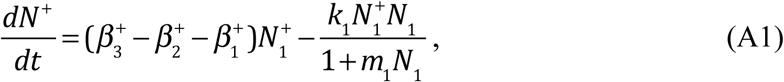

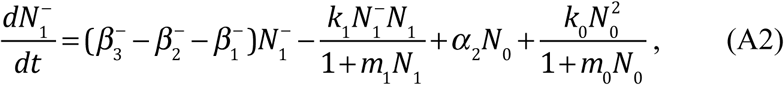

where 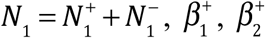 and 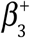 are the apoptosis, differentiation and proliferation rates in the mutant type and 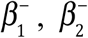 and 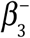 are the apoptosis, differentiation and proliferation rates in the non-mutant type. The mutant will increase in number when

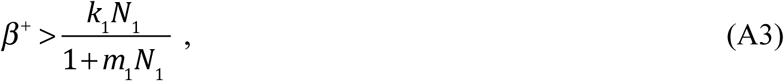

where 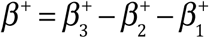, and we also define 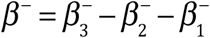. The equilibrium state for 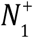, setting *dN*^+^ / *dt* = 0 in equation (A1) and assuming the mutant increases, is:

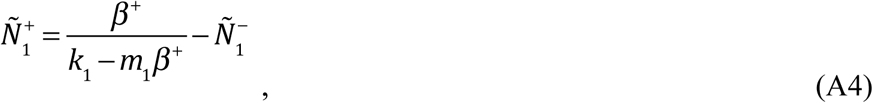

where “∼” indicates the equilibrium values. The equilibrium value for the total functional stem cell count, when the mutant spreads, is therefore

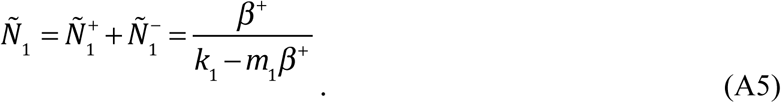

Solutions for the numbers of differentiated and semi-differentiated cell types require information on the relative numbers of mutant and non-mutant functional stem cells. The differential equation for the mutant frequency, 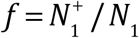, is:

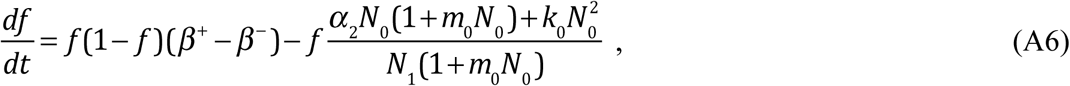

(Appendix B) which demonstrates an unusual form of natural selection that is weaker when the population size (*N*_1_) is small relative to *N*_0_ and stronger when it is large. When *N*_0_ = 0, Equation A6 reduces to a classic population genetic formula stating that the rate of allele frequency change is proportional to *f* (1− *f*), the variance in allelic states. Solving *df* / *dt* = 0 (when *β*^+^ > *β*^−^) gives the equilibrium state

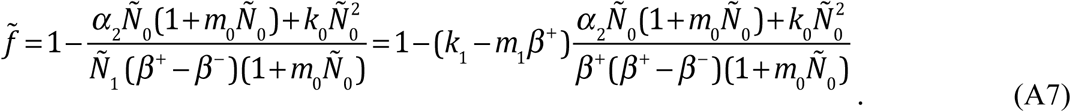

When growth is not exponential, namely not cancerous, *k*_1_ − *m*_1_*β*^+^ is positive, and so the non-mutant type is maintained via continual replenishment by the quiescent stem cells. The number of non-mutant cells is then:

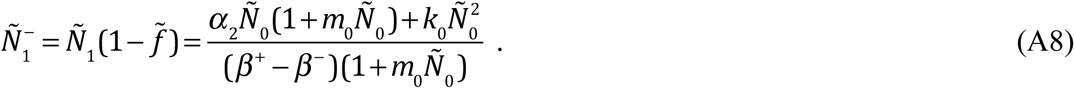

When a new mutant is introduced into a population consisting of an existing mutant type and a non-mutant type, the criterion for frequency increase is the same as for the initial mutant (Inequality A3). If increase occurs, the new mutant will entirely replace the original mutant. This is because the differential equation for the proportion of total mutants that are the new mutant type (*f*’ = *N*^++^ /(*N*^+^ + *N*^++^) where *N*^++^ is the number of new mutants), has the same form as the first term of Equation A6,

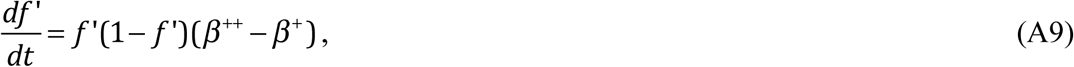

where *β*^++^ is the net growth rate of the new mutant, solving *df*’/ *dt* = 0 gives *f* = 1 when (*β*^++^ − *β*^+^) > 0 and *f* = 0 when (*β*^++^ − *β*^+^) < 0. Note that after such a replacement of a previous mutant with a new mutant type, the number of non-mutant functional cells at equilibrium will still be greater than zero (i.e. 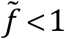, given by Equation A7) due to their active replacement by quiescent cells.

### B) Derivation of the differential equation for mutant allele frequency

The frequency of the mutant is 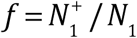, where 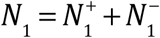. Applying the quotient rule gives:

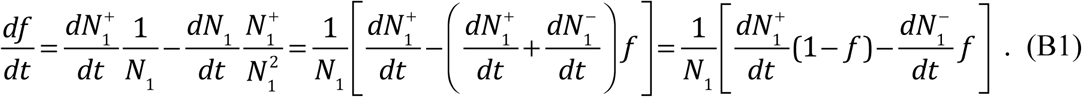

Substituting in the derivatives from Equations A1 and A2:

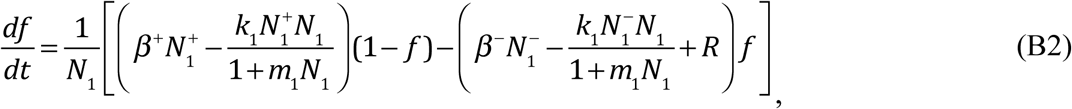

where 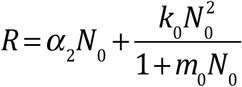. Rearranging using 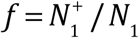 and 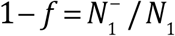 gives

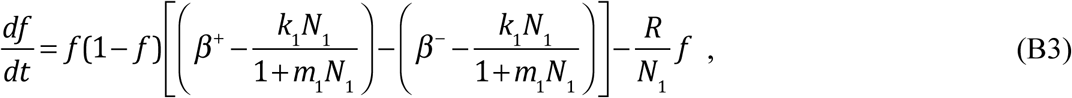

and the terms involving *k*_1_ cancel, providing the result in Equation A6:

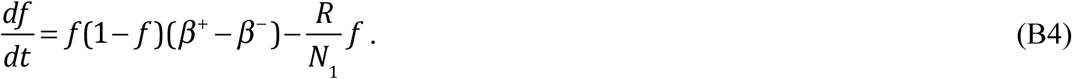

### C) Cell counts at equilibrium

The following are adopted from Johnston et al. [43, 44]:

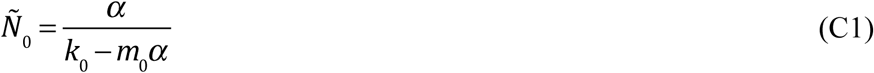

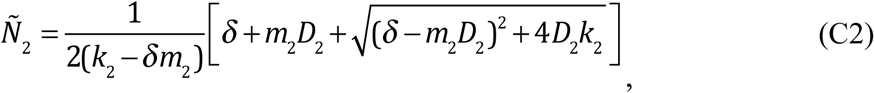

where 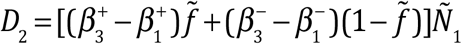, the overall differentiation rate in functional stem cells, and

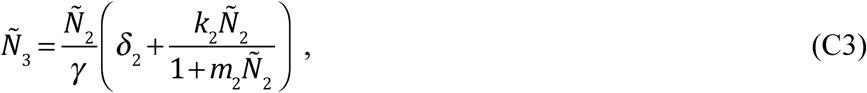

where *α* = *α*_3_ − *α*_2_ − *α*_1_ and *δ* = *δ*_3_ − *δ*_2_ − *δ*_1_. The number of semi-differentiated cells descending from mutant functional stem cells 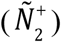, shown in Figure 2a, is given by Equation C2 replacing *D*_2_ with 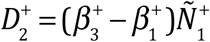 when *f* > 0, and 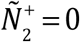 when *f* = 0.

The number descending from non-mutant functional stem cells 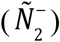 is obtained by replacing *D*_2_ with 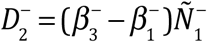. Likewise, the numbers of differentiated cells descending from mutant 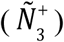 and non-mutant 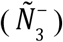 functional cells are given by Equation C3 after replacing *Ñ*_2_ with 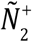 and 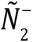 respectively.

The equilibrium state for *N*_1_ depends on whether there are mutant cells present. When there are no mutant cells at equilibrium, i.e. when all functional stem cells are genetically identical to their ancestral quiescent layer cells, the equilibrium for *N*_1_ is, from Johnston et al. [43, 44],

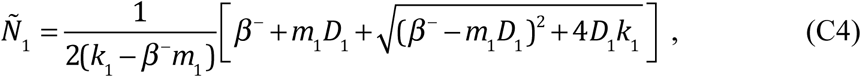

where *D*_1_ = (*α*_3_ − *α*_1_)*Ñ*_0_, the quiescent cell differentiation rate. When there are mutants that can grow fast enough to reach a stable equilibrium (Inequality A3), the stable numbers of non-mutant and mutant functional stem cells are given by Equations A4 and A8 and Equations 6 and 7 in the main text.

